# The landscape of human brain immune response in patients with severe COVID-19

**DOI:** 10.1101/2021.01.08.425999

**Authors:** John F. Fullard, Hao-Chih Lee, Georgios Voloudakis, Shengbao Suo, Zhiping Shao, Cyril Peter, Behnam Javidfar, Wen Zhang, Shan Jiang, André Corvelo, Emma Woodoff-Leith, Dushyant P. Purohit, Gabriel E. Hoffman, Schahram Akbarian, Mary Fowkes, John Crary, Guo-Cheng Yuan, Panos Roussos

## Abstract

In coronavirus disease 2019 (COVID-19), caused by severe acute respiratory syndrome coronavirus 2 (SARS-CoV-2) infection, the relationship between brain tropism, neuroinflammation and host immune response has not been well characterized. We analyzed 68,557 single-nucleus transcriptomes from three brain regions (dorsolateral prefrontal cortex, medulla oblongata and choroid plexus) and identified an increased proportion of stromal cells and monocytes in the choroid plexus of COVID-19 patients. Differential gene expression, pseudo-temporal trajectory and gene regulatory network analyses revealed microglial transcriptome perturbations, mediating a range of biological processes, including cellular activation, mobility and phagocytosis. Quantification of viral spike S1 protein and SARS-CoV-2 transcripts did not support the notion of brain tropism. Overall, our findings suggest extensive neuroinflammation in patients with acute COVID-19.

**One Sentence Summary:** Single-nucleus transcriptome analysis suggests extensive neuroinflammation in human brain tissue of patients with acute coronavirus disease 2019.

---

Coronavirus disease 2019 (COVID-19), caused by the novel severe acute respiratory syndrome coronavirus 2 (SARS-CoV-2), is currently the most urgent healthcare issue in the world. The central nervous system (CNS) is not the primary organ affected by SARS-CoV-2; however, neurological symptoms have frequently been reported in COVID-19 patients (*1, 2*). Systematically studying neurological disease in COVID-19 patients presents several challenges, including, having only a subset of the population of patients with neurological symptoms, an inability to sample CNS tissues directly, and difficulties in distinguishing direct neuroinvasion vs. systemic viremia within the brain. Brain autopsies have demonstrated acute hypoxic injury (*3*), as well as plausible SARS-CoV-2 tropism (*4-7*) in the CNS. Studies in neuronal organoids have provided conflicting results about the tropism of SARS-CoV-2 on neurons (*6, 8, 9*) or epithelial cells in the choroid plexus (*10, 11*). As such, in order to fully understand the neurological impact of COVID-19, it is critical to perform an unbiased and comprehensive high-resolution exploration of the transcriptomic landscape in human brain tissue from patients with COVID-19.

To characterize the CNS effect of SARS-CoV-2, we performed viral load quantification in human brain tissue from 5 COVID-19 patients and 4 controls (**Fig. 1a**). We note that all COVID-19 cases were classified as severe; the clinical characteristics of donors are detailed in **Data S1**. For each donor, we targeted 3 brain regions, which included the dorsolateral prefrontal cortex (PFC), medulla oblongata (medulla) and choroid plexus (ChP). Immunoblotting was negative for the presence of viral spike S1 protein in all tissues examined (**Fig. S1**). We then performed transcriptome analysis covering >99% of SARS-CoV-2 genome and all potential serotypes. For each brain region and donor, we included a single dissection with the exception of the PFC, from which we included separate dissections of cortical grey matter and white matter (for an illustrative example see **Fig. S2**), and generated, on average, 1.2 million reads per library. Across all samples, none of the sequencing reads mapped to the SARS-CoV-2 genome (**Data S2**). In addition, examination of additional brain regions (red nucleus and substantia nigra), using fluorescence *in situ* hybridization for SARS-COV-2 spike protein, also failed to detect virus (**Fig. S3**). Overall, by employing three different experimental approaches, and exploring multiple brain regions, we did not detect SARS-CoV-2 in the postmortem brain tissue.

**Fig. 1.**
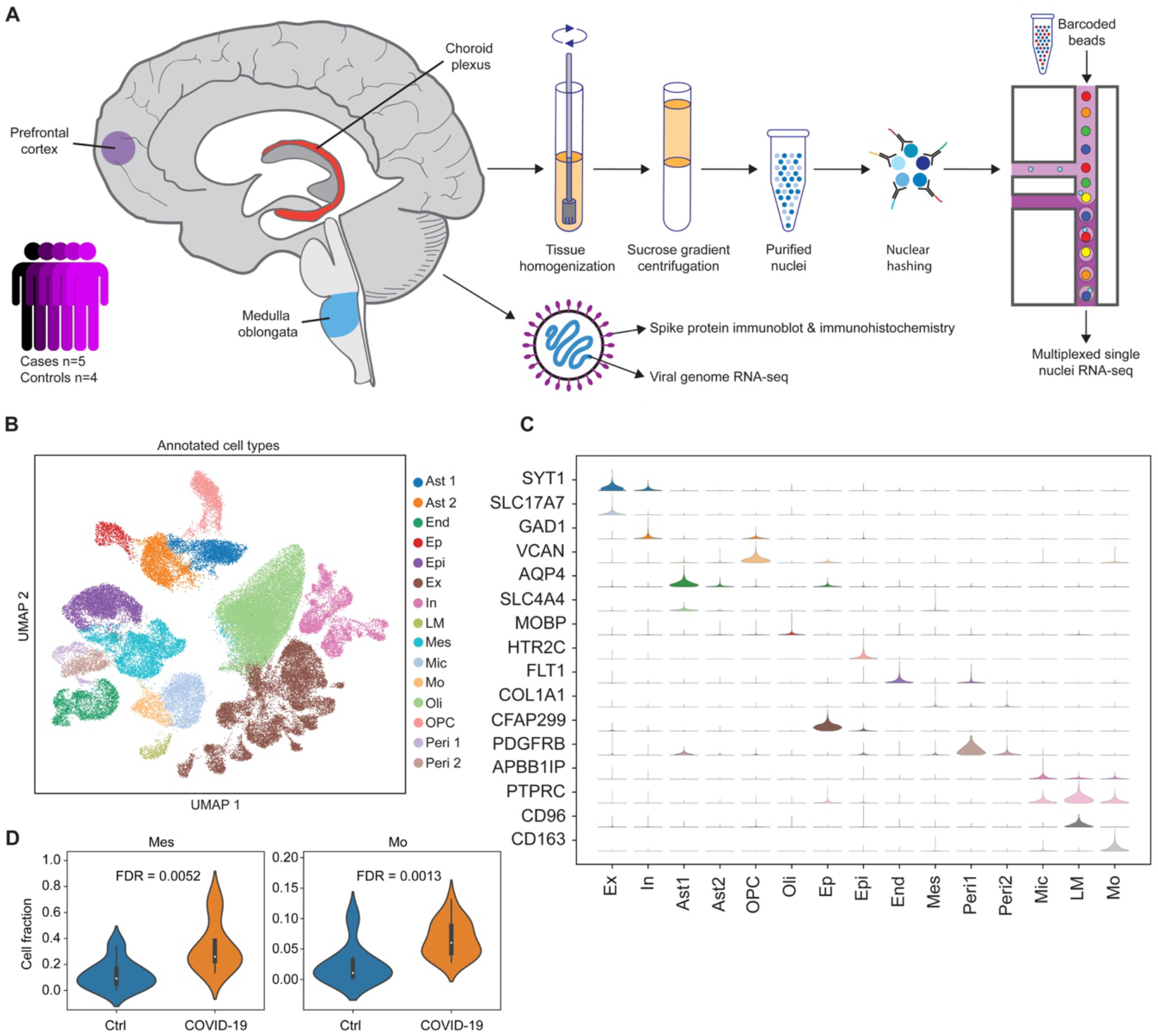
Droplet-based single-nucleus RNA sequencing in the dorsolateral prefrontal cortex (PFC), medulla oblongata (medulla) and choroid plexus (ChP) of 5 COVID-19 patients and 4 controls. **(A)**. Experimental design. Frozen specimens of human brain were dissected and subjected to a number of molecular assays, including single nuclei RNA-sequencing (snRNA-seq), viral genome RNA-seq and SARS-CoV-2 viral spike protein detection. **(B)** Uniform manifold approximation and projection visualization of annotated single nucleus data. Colors show annotated cell types. **(C)** Distribution of canonical gene markers on annotated cell populations. The range of violins are adjusted by the maximum and minimum in each row. **(D)** Cell composition of mesenchymal cells (Mes) and monocytes (Mo) in the choroid plexus stratified by case – control status. Only comparisons across tissues and cell types that survived false discovery rate correction are shown. Ast1 and Ast2: 2 groups of astrocytes. End: endothelial cells. Epi: Epithelial cells. Ep: ependymal cells. Ex: excitatory neurons. In: inhibitory neurons. LM: lymphocytes. Mes: Mesenchymal cells. Mic: microglia. Mo: monocytes. Oli: oligodendrocytes. Opc: Oligodendrocyte progenitor cell. Peri1 and Peri2: two groups of pericytes.

We then characterized the molecular and cellular perturbations in the CNS of COVID-19 patients, independent of SARS-CoV-2 direct invasion, by performing droplet-based single-nucleus RNA sequencing (snRNA-seq) in the PFC, medulla and ChP in the same set of 5 patients and 4 controls (**Fig. 1a**). To better control for donor batch effects, we performed two dissections for each brain region and individual **(Fig. S2)**, and all samples per donor (n=6) were pooled together using nuclear hashing. After preprocessing of snRNA-seq data, demultiplexing using hashtag-oligo intensity and quality control (see Methods), 68,557 high-quality single nuclei barcodes, demonstrating high technical reproducibility and a median of 2,817 genes, were available for downstream analysis (**Fig. S4**). Variance in gene expression was mostly driven by biological factors (cell type, brain regions and donor) (**Fig. S5**).

We performed *de novo* taxonomy based on the graph-based clustering and uniform manifold approximation and projection (UMAP) across all brain regions and samples, and identified 15 major cell clusters (**Fig. 1b**). Clustering was independent of donor effect and technical variables, while differences between the brain regions were preserved (**Fig. S6**). Annotation of cell clusters based on expression of canonical gene markers identified the following populations: Excitatory neurons (Ex) that express *SYT1* and *SLC17A7*; inhibitory neurons (In) that express *SYT1* and *GAD1*; Astrocytes (Ast1 and Ast2) that express *AQP4;* Ependymal cells (Ep) that express *CFAP299*; Oligodendrocyte progenitor cell (OPC) that express *VCAN*; Oligodendrocytes (Oli) that express *MOBP*; epithelial cells (Epi) that express *HTR2C*; endothelial cells (End) that express *FLT1*; Mesenchymal cells (Mes) that express *COL1A1*; Pericytes (Per1 and Per2) that express *PDGFRB*; Microglial cells (Mic) that express *APBB1IP*; lymphocyte (LM) that express *CD96* and monocytes (Mo), expressing *CD163* (**Fig. 1c; Fig. S7**). Gene set enrichment analysis showed overlap for expected molecular pathways and functions, such as myelination for oligodendrocytes, chemical synaptic transmission for excitatory and inhibitory neurons and T cell activation for lymphocytes (**Fig. S8**). The expression profiles of each cell type show high concordance with previous snRNA-seq in human brain tissue (*5, 12*), peripheral blood cells and brain organoids (*13*) (**Fig. S9**), indicating the robust definition of cell subpopulations in the current study.

We then assessed the relative proportions of the 15 major cell types in COVID-19 cases compared to controls across the three brain regions. For each cell cluster, we applied a linear mixed model to detect interaction between COVID-19 cases and brain regions, while controlling for donor effects. Among 45 combinations of cell types and brain regions, we identified 2 cell types from choroid plexus, including monocytes and mesenchymal cells, showing a significant increase in their relative proportions in COVID-19 cases (**Fig. 1d**). We did not detect any significant changes in the cell type composition of COVID-19 patients compared to controls in either the PFC or Medulla (**Fig. S10**). Overall, these results suggest that, in COVID-19, immune cells (monocytes and macrophages) extravasate from the blood vessels into the stroma of the choroid plexus, which is composed of mesenchymal cells.

For each cell type and brain region, we applied linear mixed models to identify differentially expressed genes (DEGs) among COVID-19 patients and controls, while controlling for donor effects (see Methods). Among the 15 cell types and 3 brain regions, microglia in the PFC showed the highest number of perturbations, including 178 DEGs (**Fig. 2a**). Gene set enrichment analysis identified biological processes such as “macrophage activation” (8 genes, P = 9.3×10^−6^) and “phagocytosis” (14 genes, P = 1.9×10^−6^) as being enriched with the 178 DEGs in PFC microglia (**Data S3**). To further investigate transcriptomic changes in canonical pathways, we calculated activity scores across 186 KEGG molecular pathways. We then applied linear mixed models and identified differences in average activity levels across COVID-19 patients and controls (see Methods). We identified 16 pathways showing significant differences in activity levels in microglia of the PFC (**Data S4**). As an illustrative example, we show the expression levels of the four most significant upregulated pathways (**Fig. 2b**). Together, these results suggest the strong activation of innate immune cells in COVID-19 brain parenchyma.

**Fig. 2.**
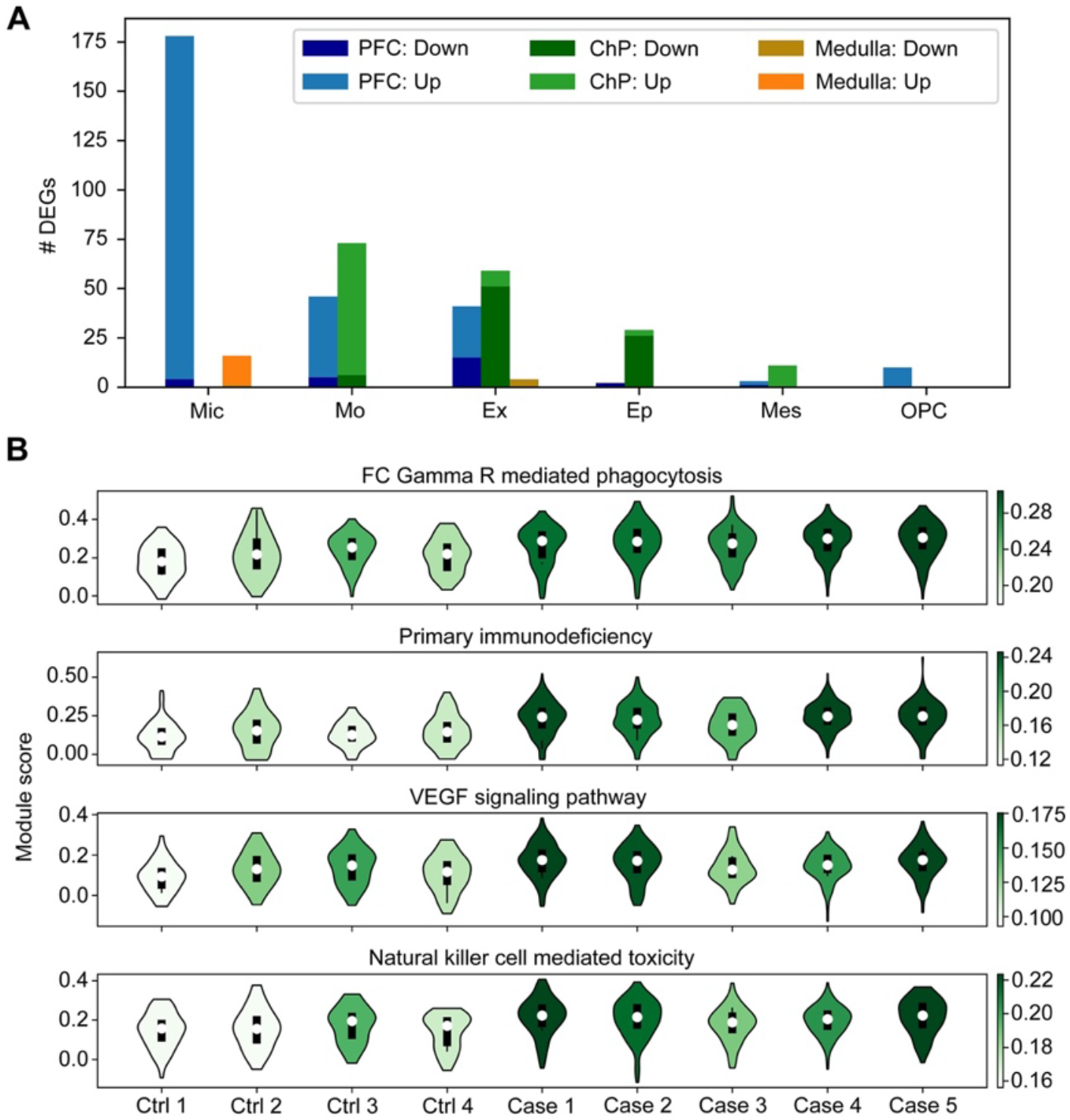
Differential gene expression and gene set enrichment analyses in COVID-19 patients compared to controls. **(A)** Number of differentially expressed genes (DEGs) identified in cell types across three brain regions. Up- and down-regulated genes are shown in different colors. Cell types are ranked by the total number of DEGs across three brain regions. Cell types with no more than 10 DEGs in any brain region are omitted. **(B)** Gene set activity scores of PFC microglia. The four most significant pathways among 186 KEGG gene sets are shown. The color of a violin indicates the median activity score of each individual.

We next calculated a pseudo-temporal trajectory score (PTS) in microglia, based on the progression of the transcriptional dysregulation in COVID-19 patients compared to controls (**Fig. 3a**). PTS has a wide distribution, potentially indicating microglia in different stages of activation and, as expected, COVID-19 patients demonstrated higher PTS than controls (**Fig. 3b**). We categorized 646 commonly expressed microglia genes into four groups (increasing, decreasing, early transient and late transient) based on the expression patterns capturing progressive changes related to PTS (**Fig. 3c, Data S5**). The majority of genes were clustered as “increasing” (579 genes), followed by late transient (36 genes), early transient (16 genes) and decreasing (15 genes). Genes within the “increasing” cluster were more perturbed in COVID-19 patients (estimated based on *pi1* = 0.683), compared to the other 3 clusters (range of *pi1* = 0.051 to 0.077) and were enriched for 452 biological pathways, including “regulation of immune system process” (136 genes, p = 1.75×10^−16^) and “apoptotic process” (104 genes, P = 3.19×10^−5^). In the “early transient” group, 8 out of the 16 genes, including CD83 and 3 heat shock proteins (HSP90AA1, HSPB1, HSPH1), belong to the “cell activation” pathway (P = 6.40×10^−6^), while the “late transient” group was enriched for the “cell mobility” pathway (P = 4.44×10^−4^).

**Fig. 3.**
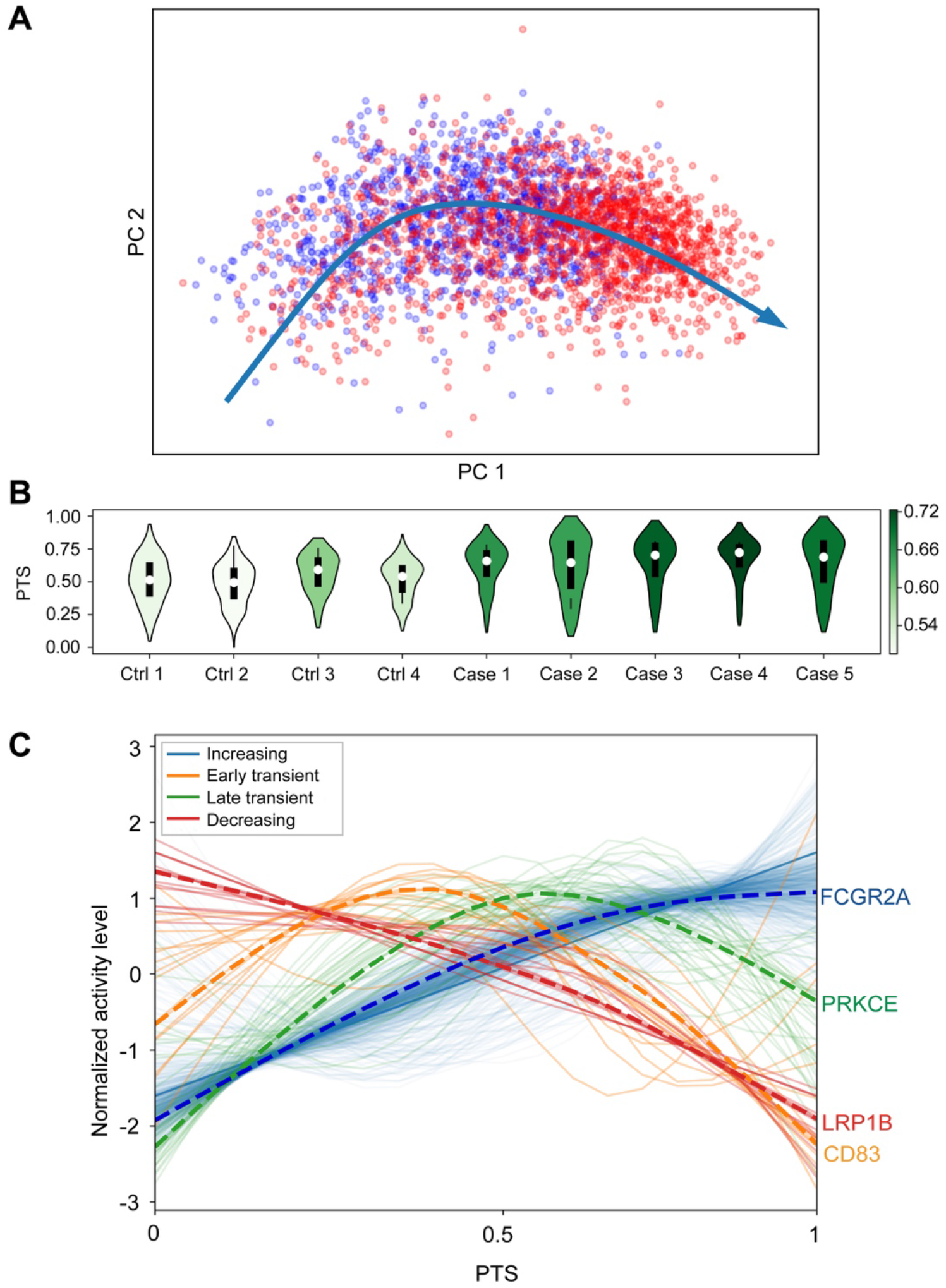
Pseudo-temporal trajectory score (PTS) analysis in microglia identified gene expression signatures with differential progression patterns in COVID-19 cases. **(A)** Pseudo-temporal trajectory in microglia across three brain regions. Red and blue colors label cells from COVID-19 cases and controls, respectively. **(B)** PTS across 5 COVID-19 patients and 4 controls in PFC microglia. The color of violin plots indicates the median activity score of each individual. **(C)** We identified 4 types of gene expression progression patterns over the pseudo-temporal trajectory: increasing (blue), early transient (orange), late transient (green) and decreasing (red). A dashed line shows the profile of a representative gene in each group.

To further understand the impact of SARS-CoV-2 on the relationships between transcription factors (TFs) and target transcripts, we explored differences in gene regulatory networks (GRNs). Across all cell subpopulations, we identified 131 TF modules that regulated, on average, 272 genes per module (*14*) (**Data S6**). UMAP projection based on activity scores of GRNs reaffirmed the robustness of annotated cell types (**Fig. S11**). Used the regulon specificity score to rank TF modules based on cell population specificity (*15*) (**Fig. S12**), we uncovered well-known cell type specific TFs, such as PAX6 for astrocytes and IRX8 for microglial cells. We then tested whether changes in the activity level of the top 5 TFs for each cell population were associated with COVID-19. PFC microglia were most affected and showed upregulation in the activity of 4 out of 5 TFs (IRF8, ATF5, SPI1, TAL1; at FDR 20%) in patients with COVID-19 (**Fig. 4a**; **Data S7**). Projecting microglia specific DEGs onto the GRNs of these 4 TFs showed the co-regulatory TF-gene patterns affected in COVID-19 (**Fig. 4b**). Collectively, these results suggest GRNs corresponding to activated microglia response in patients with COVID-19.

**Fig. 4.**
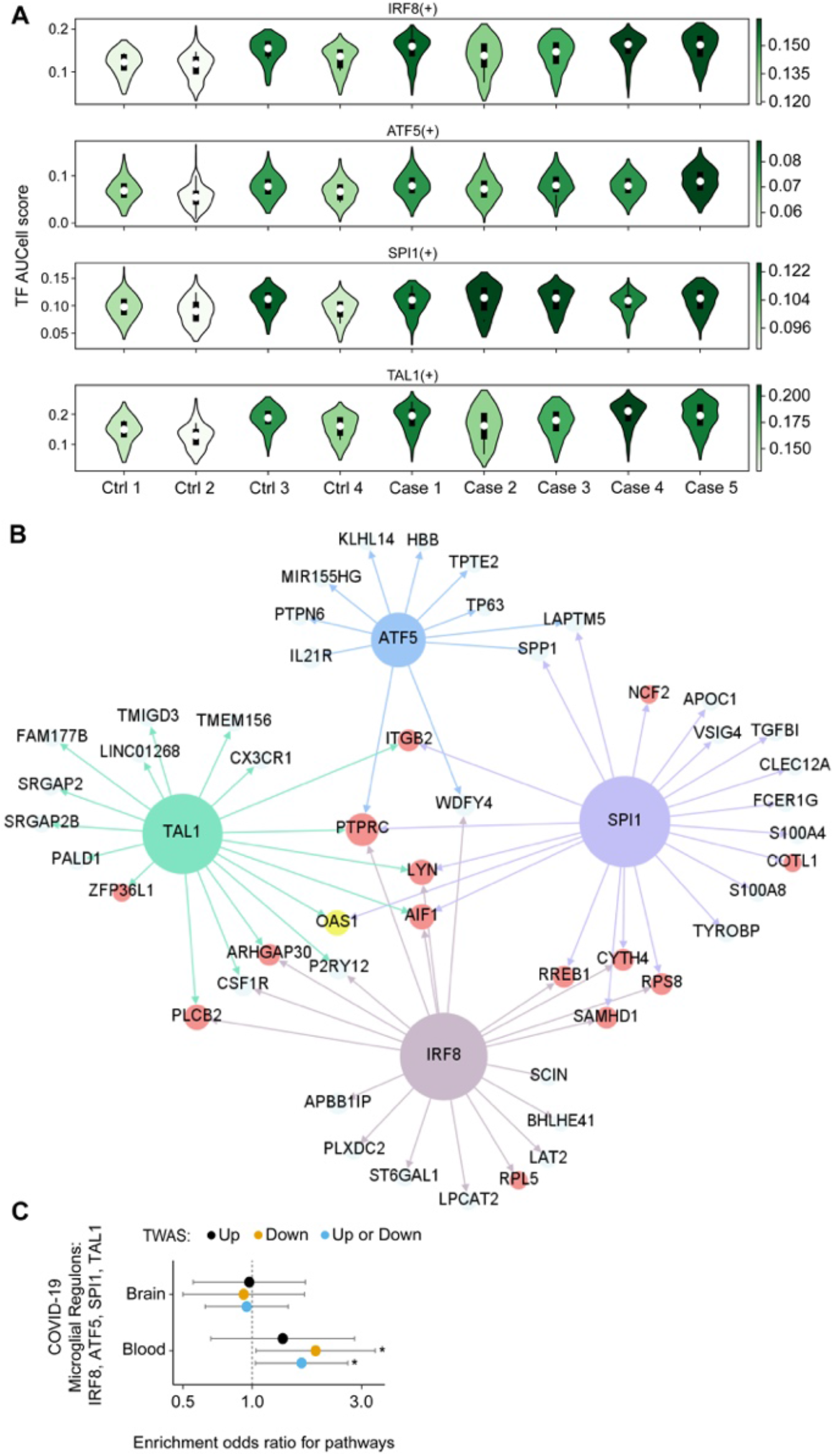
Gene regulatory network (GRN) analysis revealed transcription factors (TFs) driving transcriptomic dysregulation in COVID-19 patients. **(A)** TF module scores in PFC microglia. The color of a violin indicates the median activity score of each individual. Four up-regulated TF modules (IRF8, ATF5, SPI1, TAL1) are shown. **(B)** Up-regulated TF modules in PFC microglia. Colored nodes show the transcription factors (blue, green, brown and purple), DEGs (red) and a genetically associated gene based on GWAS (yellow). Nodes without circles are genes regulated by the transcription factors but are not DEGs. The regulatory network is trimmed to show only 14 DEGs, ranked by P-values, and 10 non-DEG genes regulated by each transcription factor. **(C)** Enrichment of the GWAS associated genes in 4 microglia specific regulons: IRF8, ATF5, SPI1, TAL1. Circles show odds ratios for the overlap of nominally significant GWAS gene (n = 285 and 560 for blood and brain respectively, P ≤ 0.05), imputed from a GWAS comparing hospitalized COVID with respect to the general population, and genes of 4 microglia specific regulons. Error bars show 95% confidence intervals of estimated odds ratios. “Up” those that are predicted to be upregulated (n = 140 and 297 for blood and brain respectively, P ≤ 0.05) and “Down” are those that are predicted to be downregulated (n = 145 and 263 for blood and brain respectively, P ≤ 0.05). Analysis is limited to protein coding genes only. Significant enrichments (P ≤ 0.05, Fisher’s exact test) are denoted by “*”.

To assess whether microglia activation has a beneficial or deleterious effect, we imputed the genetically regulated transcriptomic changes (*16*) associated with severe COVID19 outcomes by leveraging genetic variation from the COVID-19 Host Genetics Initiative and gene expression models from brain (*17, 18*) and blood (*19*) tissues (see Methods, **Data S8** for full results). We included both brain and blood tissue models to better capture the transcriptome profiling of the microglial cell lineage (*20, 21*). We identified 12 significant genes (at FDR 5%) in brain and blood (*AP000295*.*1, CCR3, CR936218*.*2, CRHR1, FYCO1, IFNAR2, IL10RB, IL10RB-AS1, LRRC37A4P, MAPT-AS1, OAS1, OAS3*) that were associated with hospitalized COVID-19 patients, with respect to the general population. Nominally significant gene-trait associations (P < 0.05) from the imputed blood transcriptome were enriched (OR=1.64, P=0.030, fisher’s exact test) with GRNs that are associated with the top 4 microglia specific TFs (IRF8, ATF5, SPI1, TAL1) (**Fig. 4c**). Enrichment was observed only for genes predicted to be downregulated (OR=1.89, P=0.035, fisher’s exact test), compared to genes predicted to be upregulated in susceptible individuals (OR=1.35, P=0.25, fisher’s exact test). In addition, *OAS1*, which was predicted to be downregulated in susceptible individuals (FDR-adjusted P = 0.003), was involved in 3 microglia specific GRNs (SPI1, IRF5, TAL1, **Fig. 4b**). Overall, the gene expression perturbations in microglia had the opposite effect compared to the genetically regulated transcriptomic changes, suggesting that microglia activation in acute COVID-19 patients represents a beneficial host response.

In summary, to better understand the impact of acute COVID-19 on the CNS, we studied its effects across 3 functionally distinct regions of the human brain (prefrontal cortex, choroid plexus and medulla oblongata). Although no virus was detected, single-nucleus gene expression analysis revealed extensive differences in brains of COVID-19 patients when compared to controls; specifically, in the ChP and PFC. We observed a relative increase in the proportions of infiltrating immune cells in the ChP, suggesting potential migration of SARS-CoV-2-carrying monocytes across the blood–brain barrier. Microglia residing in the PFC displayed dysregulated gene expression in response to SARS-CoV-2 infection. The majority of the microglial DEGs were up-regulated, mediating a myeloid-driven inflammatory response that involved a range of biological processes, including cellular activation, mobility and phagocytosis. This is consistent with previous studies (*5, 7*) that have also described increased inflammatory response of microglia in COVID-19 cases. Finally, by leveraging genetic variation to infer differences in COVID-19 susceptible individuals, we provided support for a potential beneficial role of microglia activation during the acute COVID-19 phase.

Although there is evidence that SARS-CoV-2 spike protein can be detected in the brain, including cortex (*22*), choroid plexus (*5*) and medulla oblongata (*7, 23*), immunoblotting, immunohistochemistry and viral genome RNA-seq indicate that the virus was not present at the time of death in the specimens included in this study. The ability to detect SARS-CoV-2 in the CNS is affected by the duration of COVID-19 infection (*7*). In addition, only a subset of COVID-19 patients indicates non-zero SARS-CoV-2 RNA copies in CNS, which are more difficult to detect in brain parenchyma compared to the olfactory mucosa (*7*). Although our focus was on immune cells, there is evidence, in addition to microglia activation, for COVID-19 related transcriptional changes in a range of brain cell-types including astrocytes, oligodendrocytes and excitatory neurons (*5*). The observed differences in the number of DEGs, and the cell-types affected, might be explained by the experimental design: two versus a single dissection per brain region and individual. If we only consider a single dissection per brain region and individual in our analysis, the number of DEGs increases (data not shown) and involves perturbations among every major CNS cell type.

Taken together, these findings indicate persistent activation of the innate immune response in the brains of patients with COVID-19. Based on our results, it is possible that the inflammatory response of microglia is induced from peripheral immune cells infiltrating CNS though the blood– brain barrier. Another point of entry of SARS-CoV-2 to CNS is by crossing the neural–mucosal interface in olfactory mucosa (*7*). These two mechanisms are non-mutually exclusive and might be associated with different stages of disease progression and presentation of clinical symptoms. While this study sheds light on some putative mechanisms through which SARS-CoV-2 affects the CNS, additional research is needed to deepen our understanding of the molecular mechanisms mediating neurological symptoms in COVID-19. Conclusively, our study suggests extensive neuroinflammation and brain immune response in acute COVID-19 patients, even in the absence of direct evidence of SARS-CoV-2 neuroinvasion.

## Acknowledgments

We thank the patients and families who donated material for these studies. We thank the members of the Roussos laboratory for thoughtful advice and critique and the computational resources and staff expertise provided by the Scientific Computing at the Icahn School of Medicine at Mount Sinai. This paper is dedicated to the memory of Mary Fowkes.

## Author contributions

J.F.F. and P.R. conceived of and designed the project. M.F., E.W.L., D.P.P. and J.C. provided human brain tissue and performed neuropathological examination. J.F.F., S.A. and P.R. designed experimental strategies. J.F.F. and Z.S. performed single nuclei data generation. C.P. and B.J. performed immunoblotting and fluorescence in situ hybridization. H.C.L., G.V., S.S., G.F., G.C.Y. and P.R. designed analytical strategies. H.C.L., S.S. and G.F. developed and performed all single nuclei data analyses and interpreted results. S.J. and H.C.L. performed preprocessing and quality control for all single nuclei data. A.C. preprocessed SARS-CoV-2 targeted RNA-seq data. W.Z., and G.V. developed and performed all transcriptome-wide association studies and interpreted results. J.F.F., H.C.L., G.V. and P.R. wrote the manuscript with the help of all authors. P.R. supervised overall data generation and analysis.

## Supplementary Materials

Materials and Methods Figs. S1 to S12

Captions for Data S1 to S8 References

