## Supplement for "The landscape of human brain immune response in patients with severe COVID-19"

**This PDF file includes:**

Materials and Methods
Figs. S1 to S12
Captions for Data S1 to S8
References

**Other Supplementary Materials for this manuscript include the following:**

Data S1 to S8 (Excel files)

### Materials and Methods

Brain tissue dissections

Brain tissue was harvested based on a modification of published protocols (*1*) for rapid dissection and systematic neuroanatomical sampling. First, the myelencephalon was removed, sectioned horizontally at 0.2 cm intervals with alternating sections fixed in 10% neutral buffered formalin or frozen at -80°C. Then the remaining specimen was bisected sagittally, and half fixed in 10% neutral buffered formalin. The mesencephalon and metencephalon were removed, sectioned horizontally, parcellated, barcoded and rapidly frozen between two metal plates previously cooled to -80°C. The forebrain was then sectioned coronally at ~0.4 cm and systematically dissected, barcoded and frozen at -80°C as well. For each donor, we targeted 3 brain regions, which included the dorsolateral prefrontal cortex (PFC), medulla oblongata (medulla) and choroid plexus (ChP). To better control for donor batch effects in the snRNA-seq analyses, we performed two dissections for each brain region and individual (for an illustrative example see **Fig. S2**). For the immunoblotting and SARS-CoV-2 targeted RNA-seq assay, we included a single dissection with the exception of the PFC, from which we included separate dissections of cortical grey matter and white matter.

Western Blot Analysis of COVID-19 Receptors in Human Brain

100mg (1-volume) of brain tissue was Dounce homogenized in 1-ml (10 volumes) of ice-cold protein extraction buffer (50 mM-Tris-HCl, pH 8; 20 mM-NaCl; 2 mM-MgCl2; 4M-Urea, 0.35% Triton-X100; 0.35% Sodium deoxycholate; 0.05% SDS; supplemented with protease and phosphatase inhibitors). Benzonase was added to eliminate viscosity of the sample for 10 min on ice, then sonicated and centrifuged at 20,817 g for 15 minutes at 4°C. Clarified total lysates were quantified using a BCA Protein Assay kit (Pierce). Western blot analysis was performed using 100μg of total protein using Anti-SARS-CoV-2 spike glycoprotein (Abcam, ab272504, 1:5,000) or anti-β-actin (Abcam, Ab4970S) primary antibodies (1:5000, 1-hr) followed by autoradiographic detection using corresponding anti-rabbit (Cell Signaling, Cat#7074S) secondary antibodies conjugated with HRP (1:5000, 1-hr). Purified SARS-CoV-2 (COVID-19) S1 Recombinant Protein (ProSci Inc, Cat#97-086) was used as positive control to track the presence COVID-19 in brain samples.

SARS-CoV-2 targeted RNA-seq

RNA was extracted from each individual dissection using the miRNeasy kit (Qiagen, Cat no: 217004), according to manufacturer's instructions. RNA samples were shipped to the New York Genome Center and went through an initial quality check. For the SARS-CoV-2 targeted assay we used the AmpliSeq Library Plus and cDNA Synthesis for Illumina kits (Illumina, Cat nos: 20019103 & 20022654). Briefly, 20ng of RNA was reverse transcribed, the cDNA targets were then amplified with the Illumina SARS-CoV-2 research panel (Illumina, 20020496). The amplicons were partially digested and AmpliSeq CD Indexes were ligated onto the amplicons. The library was then cleaned up and amplified. After amplification there was a final clean up and the libraries were quantified, pooled and run on a NovaSeq 6000 S4 in a 2x150 run format.

SARS-CoV-2 genome consensus calling and taxonomic classification

Short read-data were filtered and processed prior to alignment. Read pairs that did not contain a single 19bp seed *k*-mer in common with the SARS-CoV-2 genome reference (NC_045512.2) were discarded. Adapter sequences and low quality (Q < 20) bases were trimmed from the remaining reads, using Cutadapt v2.10 (*2*). Processed reads were then mapped to the SARS-CoV-2 genome reference using BWA-MEM v0.7.17 (*3*) and only read pairs with at least one alignment spanning a minimum of 42 bp in the reference and starting before position 29,862 (to exclude polyadenine-only alignments) were kept and used for genome consensus calling. Additionally, all short-read data were taxonomically classified using taxMaps (*4*). As part of the taxMaps pipeline, reads were processed prior to mapping. Adapter sequences and low quality (Q < 20) bases were trimmed out and low complexity reads discarded. The remaining reads were then concurrently mapped against 1) the phiX174 reference genome (NC_001422.1); 2) the SARS-CoV-2 reference genome (NC_045512.2); and 3) a combined index encompassing the entire NCBI's nt database, RefSeq archaeal, bacterial, fungal, protozoan and viral genomes, as well as a selection of RefSeq model organism genomes, including the human GRCh38 reference (*5*), to produce the final classification.

RNA FISH

Human ventral midbrain sections were cut 10um on a cryostat and mounted on Superfrost Plus slides (Fisher). The slides were incubated in cold 4% formaldehyde for 10 minutes, followed by an ethanol series wash. Slides were either used immediately or stored at -20°C in 100% ethanol for up to one week. Sections were dried at room temperature to remove ethanol before being processed per the RNAscope Multiplex Fluorescent v2 protocol (Advanced Cell Diagnostics). Briefly, sections were treated with hydrogen peroxide, protease treated and incubated with anti-sense DNA probes against the SARS-CoV-2 spike protein RNA sequence for 2 hours at 40°C. Amplifier sequences were polymerized to the probes and treated with horseradish peroxidase and Opal 570 dye. Prior to mounting in VECTASHIELD, the slides were incubated for 30 seconds in TrueBlack Lipofuscin Autofluorescence Quencher (Biotium) and rinsed with 1x PBS. Images were acquired at 20x and 63x on a Zeiss LSM780 Confocal microscope.

Tissue processing, nuclei hashing and multiplexed single nuclei RNA-seq

RNase inhibitors (Takara) were used throughout. 50 mg of frozen human tissue was homogenized in cold lysis buffer (0.32 M Sucrose, 5 mM $CaCl_2$, 3 mM Magnesium acetate, 0.1 mM, EDTA, 10 mM Tris-HCl, pH8, 1 mM DTT, 0.1% Triton X-100) and filtered through a 40μm cell strainer. The flow-through was under-laid with sucrose solution (1.8 M Sucrose, 3 mM Magnesium acetate, 1 mM DTT, 10 mM Tris-HCl, pH8) and centrifuged at 107,000 g for 1 hour at 4°C. Pellets were re-suspended in 0.5 ml DPBS. Following re-suspension nuclei were quantified using a Countess automated cell counter (Life technologies). 6 tissue samples were prepared in parallel and 2M nuclei from each were set aside and incubated with individual nuclear hashing antibodies (TotalSeqA anti-Nuclear Pore Complex Proteins, BioLegend) according to manufacturer's instructions, to facilitate multiplexed loading of the single-cell micro-fluidics device (10x Genomics). Following antibody incubation, nuclei were washed twice, quantified, normalized to 1000/μl and pooled. A total of 46,560 nuclei (7760 each) were loaded, in duplicate, on 10x Genomics B chips using 3' capture chemistry. snRNA-seq library construction was performed according to manufacturer's instructions (10x genomics). In parallel, hash-tag oligos (HTOs) were generated separately, as previously described (*6*). Prior to sequencing, library concentration and molecular weights were determined by QPCR (KAPA biosystems) and Tapestation (Agilent Technologies), respectively. Finally, libraries were sequenced on the Nova-seq platform (Illumina) obtaining 2x100 paired-end reads.

Single nucleus data processing and demultiplexing

Paired-end sequencing reads were processed using cell ranger v3.1.0. We aligned sequencing reads to a pre-mRNA reference genome to capture un-spliced pre-mRNA, as recommended by 10x genomics. We used kallisto indexing and tag extraction (KITE) to quantify HTO barcodes (*7*). 6 HTO tags, corresponding to two repeated dissections of three brain regions, were quantified for each sample.

We customized an algorithm to identify singlets and doublets using HTO barcodes. Briefly, the HTO barcodes were first grouped into 6 clusters using k-means clustering for every sample. For each HTO associated with a dissection, we calculated a cut-off to classify cells into HTO positives and negatives. The cut-off is determined by fitting a logistic regression model to classify the cluster with highest average HTO counts against the cluster with lowest average HTO counts. After thresholding, cells that are positive in zero, one or two HTO barcodes are assigned as negatives, singlets, and doublets. We iterated the process by classifying singlets against other cells to refine the final classification.

Demultiplexed UMI matrices across independent experiments were combined and labeled by their associated experiment batch and brain region. Nuclei with fewer than 200 expressing genes or higher than 5% mitochondrial reads were omitted. We applied DoubletFinder (*8*) to remove doublets that were missed by HTO demultiplexing. We used Seurat (*9*) to perform data normalization, UMAP dimension reduction and graph-based clustering. We first normalized the count matrix to a uniform coverage of 10,000 reads followed by log transformation. Transformed values were capped by 10. Ten thousand variable genes were identified using variance stabilizing transformation implemented in Seurat FindVariableFeatures function. Harmony (*10*) was used to adjust batch effects using the first 50 principal components of the transformed count matrix. UMAP projection and Louvain clustering were then performed based on the shared nearest neighbor graph of these 50 harmonized principal components. A preliminary inspection on clustering and UMAP projection identified a cluster of doublet cells and a cluster of cells that are potentially introduced due to dissection bias. These two clusters were omitted from the downstream analysis. We then repeated the process to generate the final clustering and UMAP projection. We annotated clusters by inspecting a selected set of canonical markers.

Statistical analysis on cell composition

To identify cell composition changes associated with COVID-19, we computed the fraction of cells annotated with a cell type in each brain region and then fitted a linear mixed model:

$$\text{cell fraction} \sim \text{COVID-19} \mid \text{tissue} + (1 \mid \text{tissue}) + (1 \mid \text{donor}).$$

Associated cell types were then identified by contrasting cases and controls for each tissue. We repeated statistical tests on the normalized cell fraction using centered log ratio transformation and obtained similar results.

Identification of differentially expressed genes

Differentially expressed genes (DEGs) were identified using a linear mixed model. We first split the combined count matrix of 10,000 variable genes into sub-matrices containing nuclei from one annotated cell population. K-nearest neighbor smoothing (*11*) was applied to each sub-matrix to impute dropped-out genes. Genes expressed in less than 1% cells in a cell population were further filtered. A pseudo bulk count matrix was formed by aggregating single cell reads per sample. Using the dream pipeline (*12*), we fitted a mixed linear model:

$$\text{Log( cpm( Gene expression ) )} \sim \text{COVID-19} \mid \text{tissue} + (1 \mid \text{tissue}) + (1 \mid \text{donor})$$

to the pseudo bulk count matrix. DEGs were then identified by contrasting cases and controls for each tissue. We omitted test results on genes that were sparsely expressed (<1% cells) in each tissue.

Cell type enrichment

We performed Fisher's exact test to compare gene signatures of annotated cell types with those reported in other studies. To generate the set of marker genes for an annotated cell type, we used Wilcoxon Rank Sum test, implemented by the FindMarker function, to compare cells of one cell type against the rest. Up to 50 marker genes, with adjusted p-value less than 0.05 and ranked by fold changes, were included to form the set of marker genes.

We curated gene sets of known cell types from 4 publicly available data sets:

1. Gene sets reported by (*13*), including annotated cell types in the cortex. We filtered the reported marker genes to ensure the log fold changes are larger than 0.25.
2. Gene sets obtained using the single cell data from human organoids (*14*). We used the FindMarker function to compute the marker genes for the cell type annotated as in the original study. Genes with adjusted p-value less than 0.05 and log fold change larger than 0.25 were included.
3. Gene sets obtained using the single cell data from choroid plexus (*15*). We used the FindMarker function to compute the marker genes for the cell type annotated as in the original study. Genes with adjusted p-value less than 0.05 and log fold change larger than 0.69 were included.
4. Gene sets computed using peripheral blood samples from 10x genomics. The data set includes single cell RNA-seq of 6 cell types sorted by FACS. We computed marker gene sets using the FindMarker function. Genes with adjusted p-value less than 0.05 and log fold change larger than 0.25 were included.

The background gene sets were set to be the intersection of 10,000 variable genes and all possible genes from the other data source.

Construction and analysis of the transcription factor-gene network

We used pySENIC (*16, 17*) to identify transcription factor regulons. The count matrix of 10,000 variable genes was used. Genes expressed in less than 1% cells were further filtered as recommended by the pySCENIC protocol. The gene co-expression network was inferred using the gradient boosting machine implemented by arboreto. Enriched motifs for a gene co-expressing module were predicted using pre-computed databases from cisTargetDB and the ctx function in pySCENIC. Lastly, activity scores of inferred regulons were quantified at the single-cell level using AUCell.

To identify regulons associated with COVID-19, we first computed the averaged AUCell score per sample and fitted a linear mixed model

$$\text{regulon activity} \sim \text{COVID-19} \mid \text{tissue} + (1 \mid \text{tissue}) + (1 \mid \text{donor}).$$

Associated regulons were then identified by contrasting cases and controls for each tissue. We restricted our analysis to the 5 regulons that are most specific to a cell type ranked by the regulon specific score (*18*).

Pseudo-temporal trajectory score

To construct a pseudo time trajectory that captures disease progression in microglia, we computed the principal components using the gene count matrix of the top 500 DEGs across three brain regions. Genes expressing in less than 1% cells were excluded. The resulting gene matrix includes 130 genes, accounting for duplicated DEGs that are significant in more than one brain region, and 3,052 microglia barcodes. The gene matrix was normalized to a uniform coverage of 10,000 reads and log transformed. Principal component analysis was performed on this transformed gene matrix. Five clusters were identified from the first 50 principal components using the Leiden algorithm. We then constructed a pseudo time trajectory along the first two principal components using Slingshot (*19*). The order of the pseudo time trajectory is chosen so that the fraction of cells from COVID patient is increasing with the pseudo time.
We then analyzed gene expression profiles over the pseudo time trajectory using tradeSeq (*20*). We restricted our analysis on 690 genes expressing in more than 50% of cells to mitigate the issue of zero inflation. The generalized additive model (GAM) with 6 knots was fitted to each gene. By testing if the coefficients of a GAM are constant over time (*20*), we found that 646 genes were associated with the pseudo time progression, of which 15 and 320 are strictly decreasing or increasing, respectively. Applying resampling-based sequential ensemble clustering to genes with non-monotonic profiles, we identified 22 clusters but a strong similarity between clusters was observed. We performed another clustering on the average profiles of these 22 clusters, sampled by 20 points equally spaced over pseudo time. We used dynamical time warping (DTW) to measure the similarity between average profiles. Spectral embedding based on the DTW distance suggested that the average profiles of these 22 clusters can be arranged based on the timing when expression reaches its extremum. We then grouped these 22 clusters into 6 meta-clusters using KMeans clustering and further merged 6 meta-clusters into 4 categories after manual inspection.

Transcriptome-wide association study and geneset enrichment

For the COVID-19 TWAS we used the "B2_ALL_eur_leave_23andme" summary statistics from the Release 4 (October 2020) of "The COVID-19 Host Genetics Initiative" (6,406 cases and 902,088 controls; retrieved from https://www.covid19hg.org/) which corresponds to the phenotype "Hospitalized covid vs. population, leave out 23andMe". We employ blood and brain EpiXcan (*21*) tissue imputation models. For the blood, we used the blood imputation model from the STARNET cohort as previously described (*21*), which is based on a previously published study (*22*). For the brain, we used genetic and gene expression data from PsychENCODE (*23, 24*) and generated the EpiXcan imputation model as previously described (*21*). For the generation of all imputation models (which are currently limited to samples of European origin): a) genotype datasets were uniformly processed for quality control and missing variants were imputed using the University of Michigan server (*25*) with the Haplotype Reference Consortium (HRC) reference panel (*26*), b) RNA-seq gene level counts were adjusted for known and hidden confounds, followed by quantile normalization and the residualized counts were used in the regression models. To derive gene-tissue-trait associations, transcriptomic imputation models were applied to the summary statistics following the S-PrediXcan approach (*27*). For the transcriptomic imputation, genetic variants to which any of the following apply were excluded: minor allele frequency higher or equal than 0.01, insertions/deletions, strand-ambiguous SNPs, variants that share coordinates with other variants, and variants in the broad major histocompatibility complex (MHC) region (chromosome 6: 25–35 Mb). For each tissue, we kept genes with "pred_perf_r2" value (*28*) ($r^2$ of the correlation between cross-validated prediction and observed expression) greater than, or equal to, 0.01. For the geneset enrichment analysis, only protein coding genes were considered and the enrichment was tested with the Fisher's exact test.

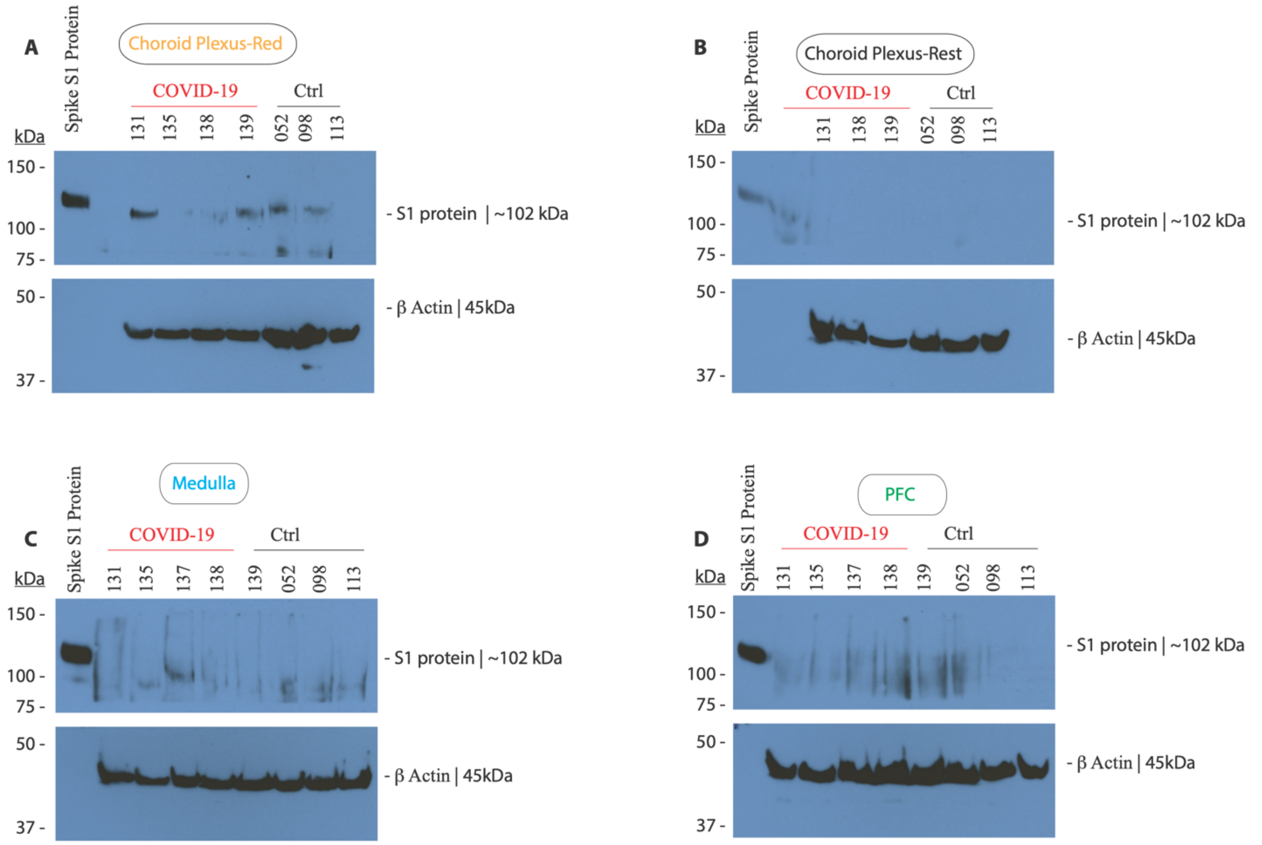


**Fig. S1. Western Blot Analysis Investigating the presence of COVID-19 in Human Brain.** Total Protein (100 µg) from infected (in red) or control brain domains including Choroid Plexus (A & B), Prefrontal cortex (D) and Medulla (C) were probed with antibody against viral spike S1 protein (102 kDa) (top blot, all panels). Purified recombinant S1 protein (5 µg/lane) was used as positive control (top, leftmost lane, all panels); Sample loading was verified using β-Actin (45 kDa) antibody (bottom blot, all panels)

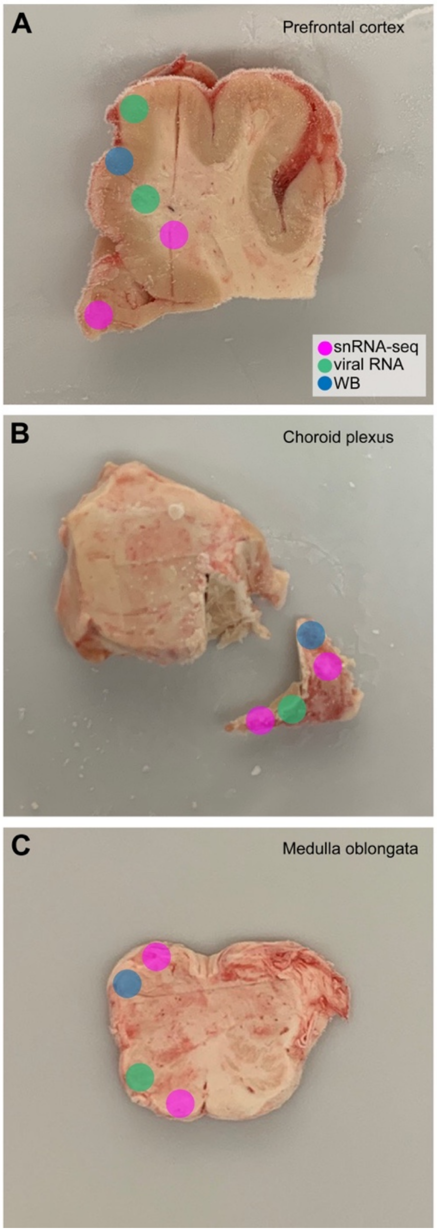


**Fig. S2. Brain specimen dissections.** Frozen brain specimens were further dissected and subjected to snRNA-seq (magenta), viral genome RNA-seq (green) and viral spike S1 protein immunoblotting (blue). (A) prefrontal cortex (PFC); snRNA-seq and viral genome RNA-seq were performed in separate dissections of cortical grey matter and white matter, (B) choroid plexus and (C) medulla oblongata.

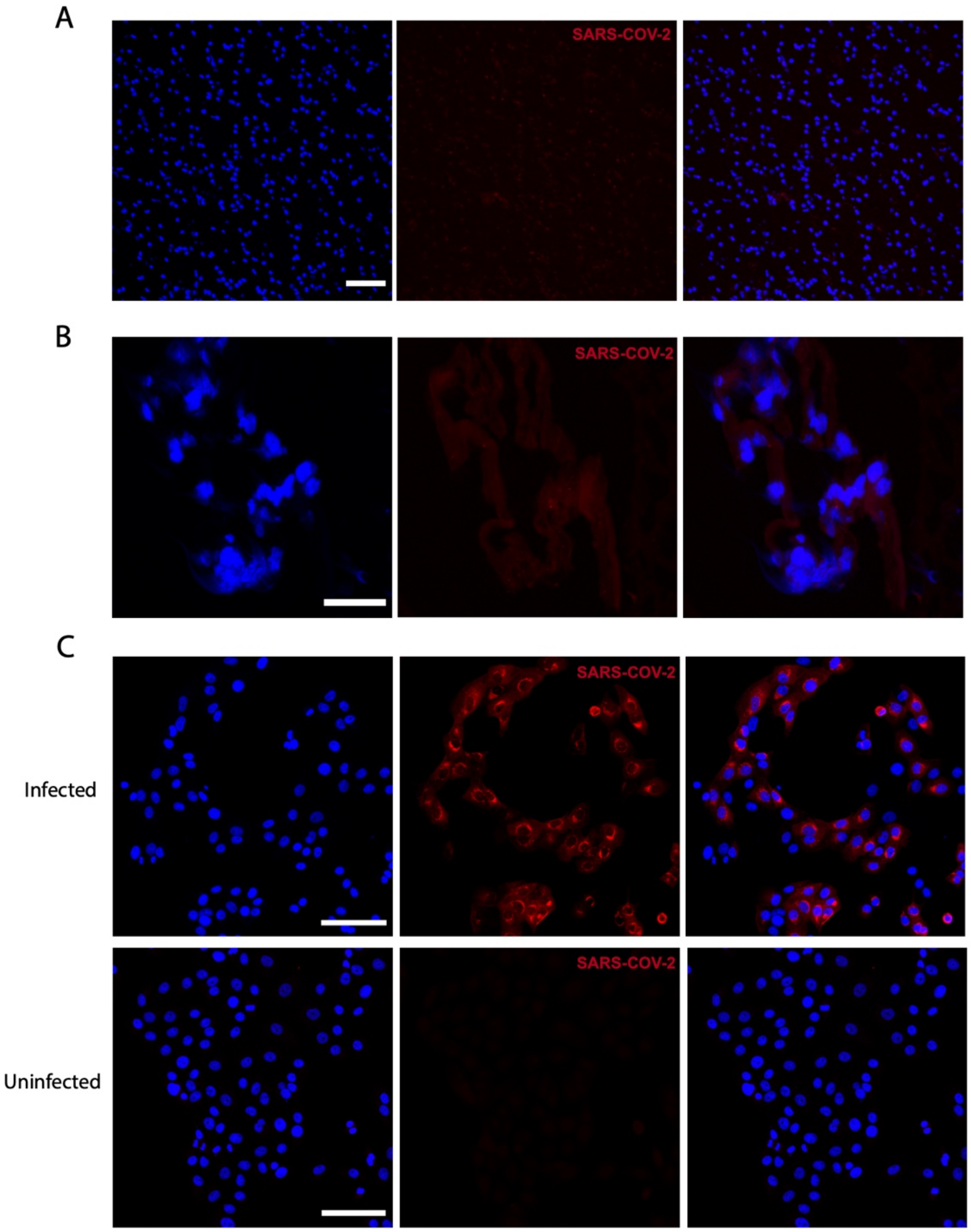


**Fig. S3. SARS-COV2 RNA not detected in postmortem midbrain of COVID-19 patients.** Fluorescence in situ hybridization for anti-sense RNA did not reveal any neurological invasion of SARS-COV-2 in the red nucleus (A) or substantia nigra (B). Midbrain blood vessels were also negative (B). A VERO E6 cell line infected with SARS-COV-2 showed robust staining for viral RNA (C). Scale bars = 100 µm (A, C); 50 µm (B).

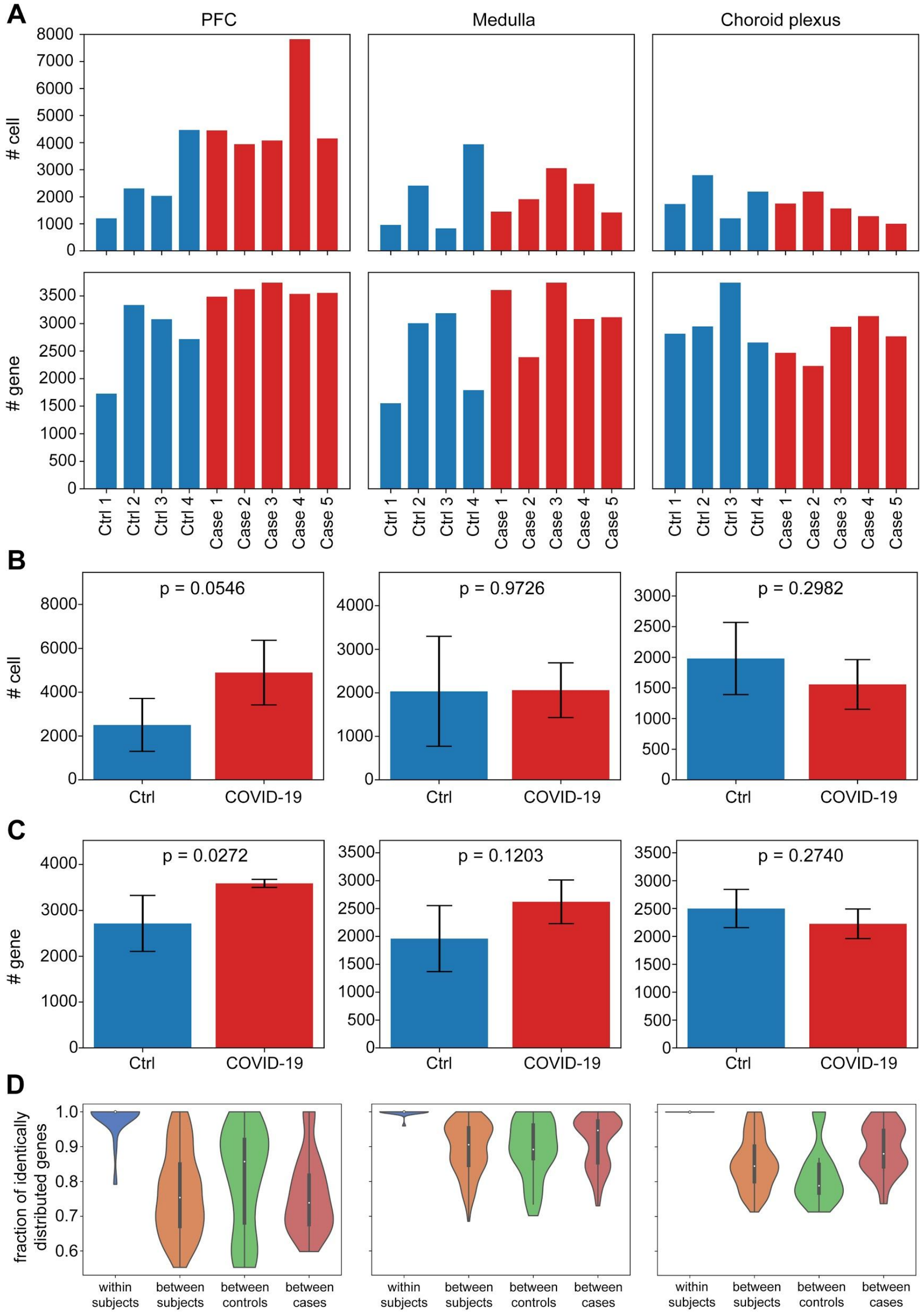


**Fig. S4. Quality control of single nucleus data.** (A) number of nuclei and expressing genes in each sample. Average numbers of (B) nuclei and (C) expressing genes. Error bars show standard deviation. P values tested by two-sample t tests are shown in each panel. (D) Number of genes that have the same distributions between two samples. Comparisons were made using two replicated sequencing samples from the same dissection (within subjects) and sequencing samples from two different subjects (between subjects), two control subjects (between contrls) and two COVID-19 patients (between cases). We tested if a gene is identifiably distributed between two samples by Wisconsin rank sum test and reported the number of genes with false discovery rate above 0.05.

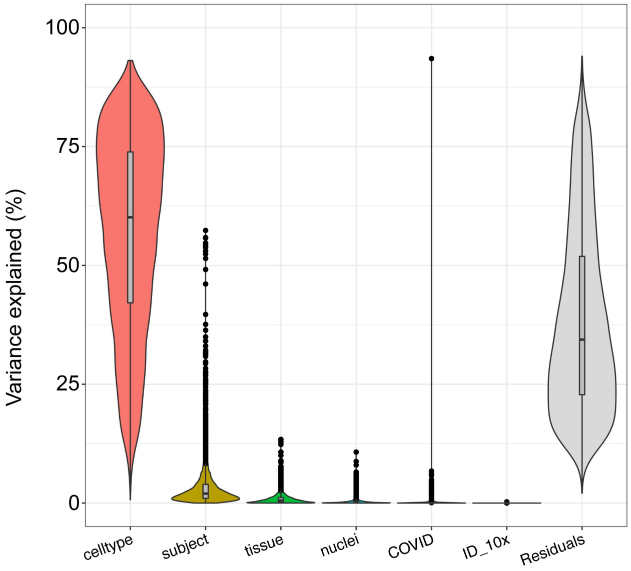


**Fig. S5. Variance partition plot.** The variance partition decomposes variance in gene expression into different covariates, including (in decreasing order for variance explained): cell type (celltype), donor (subject), brain region (tissue), number of nuclei detected in each snRNA-seq library (nuclei), COVID19 positive status (COVID) and 10x Genomics lane (ID_10x). The fraction of unexplained variance is included as "Residuals".

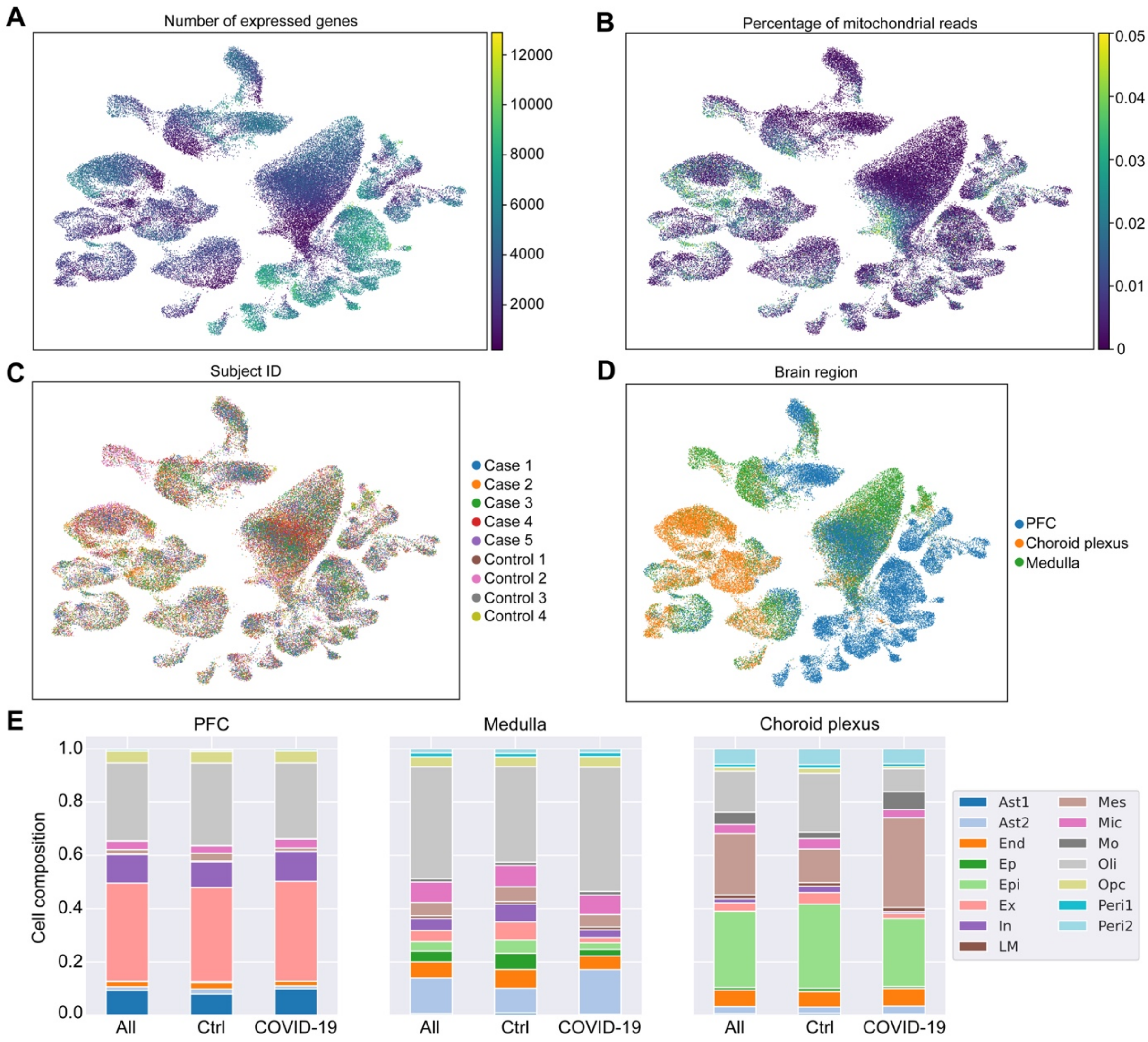


**Fig. S6. Cell type composition across 15 major cell clusters.** UMAP visualization demonstrates the distribution of (A) number of expressed genes, (B) percentage of mitochondrial reads and the origin of cells from (C) donors and (D) brain regions. (E) Overall cell composition in three brain regions. Cell composition were calculated using all samples (All), control subjects only (Ctrl) and COVID-19 patients only (COVID-19).

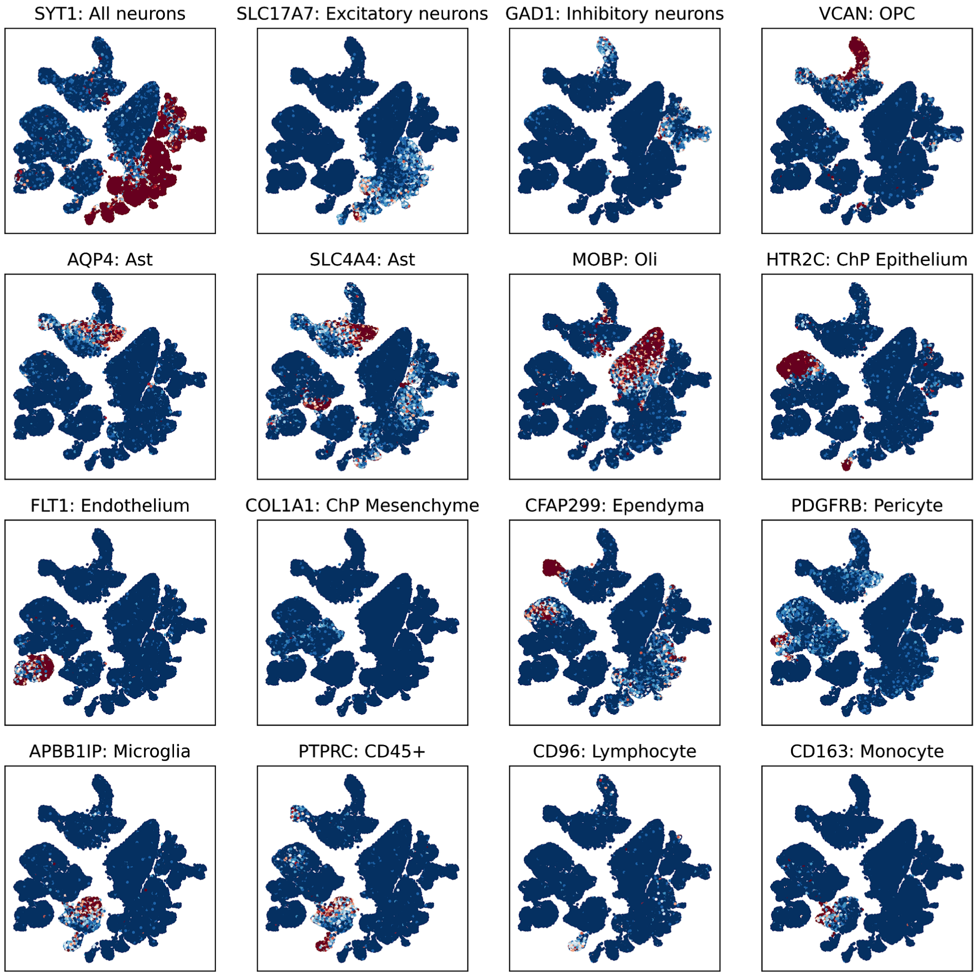


**Fig. S7. UMAP visualization of the distribution of canonical gene markers on annotated cell populations.** Each plot visualizes a single marker. Markers and corresponding cell type are described at the top of each plot.

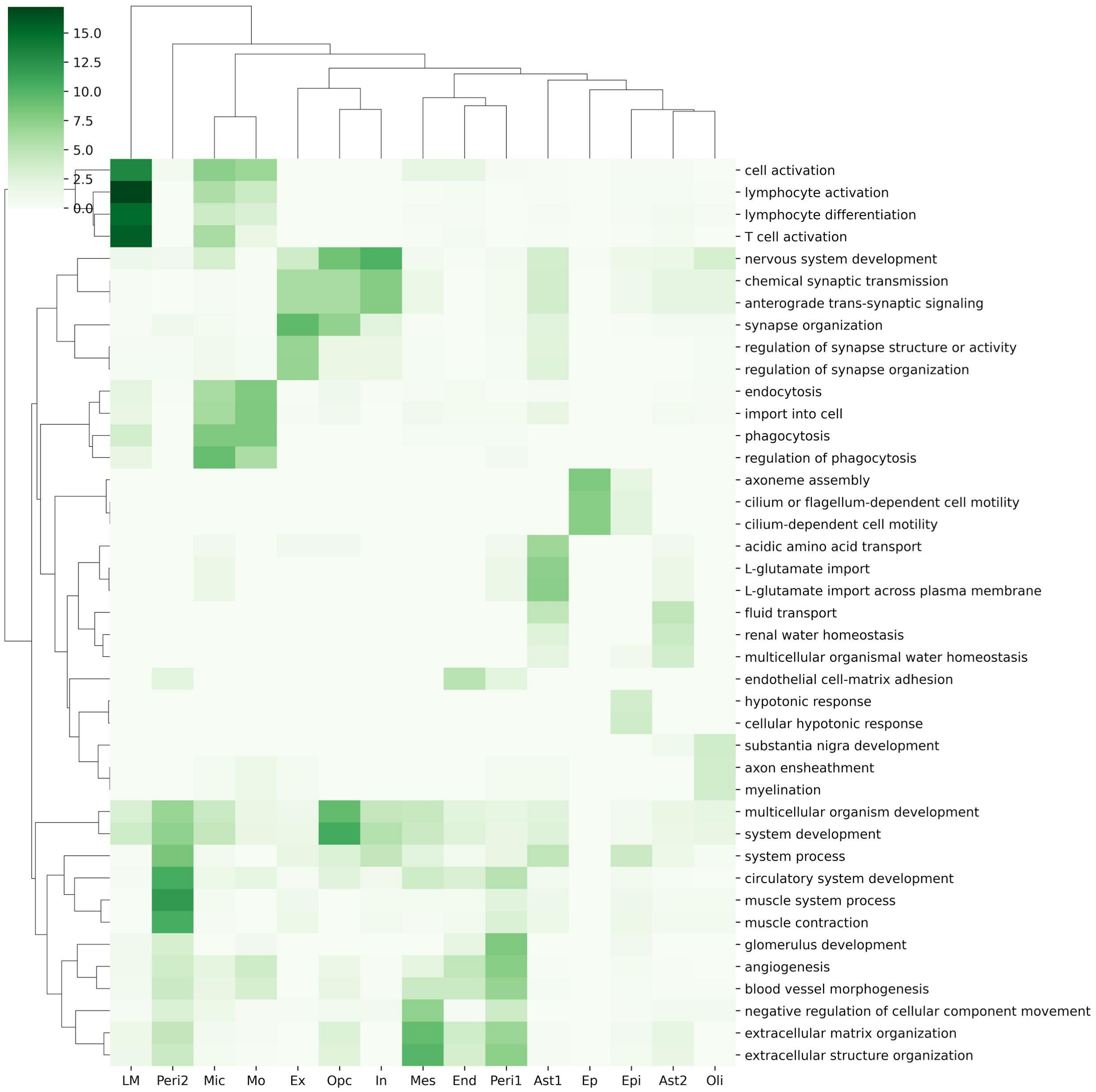


**Fig. S8. Gene set enrichment analysis for marker genes of annotated populations.** Shade of Green indicates the negative logarithm of p values of overlaps between annotated cell type (columns) and Gene Ontology terms (rows). P values were calculated by Fisher's exact test.

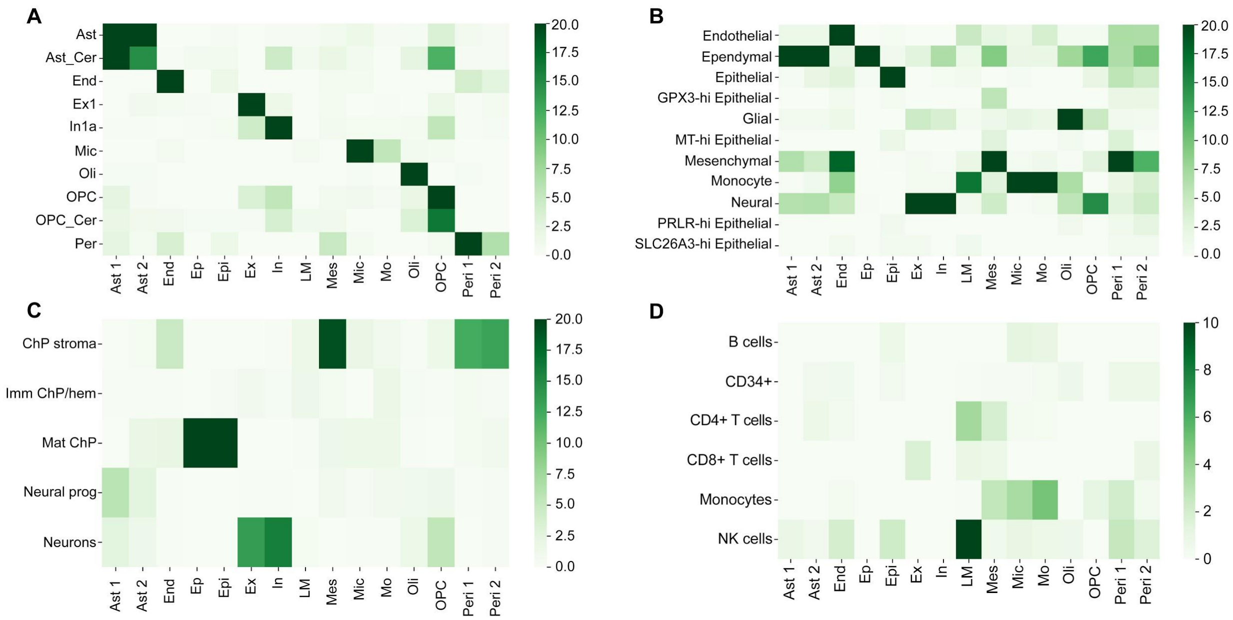


**Fig. S9. Comparison of marker genes of annotated populations from the current study and other published studies**. (A) Signatures determined from single nucleus RNA-seq data from the cortex (*13*). (B) Signatures determined single cell RNA-seq data from human central nervous system organoids (*14*). (C) Signatures determined from single nucleus RNA-seq data from choroid plexus (*15*). (D) Signatures determined from single cell RNA-seq data from peripheral blood cells sorted by fluorescence activated cell sorting (from 10X Genomics). Shade of green shows the negative logarithm of p values comparing signatures from other studies (columns) and from this work (rows).

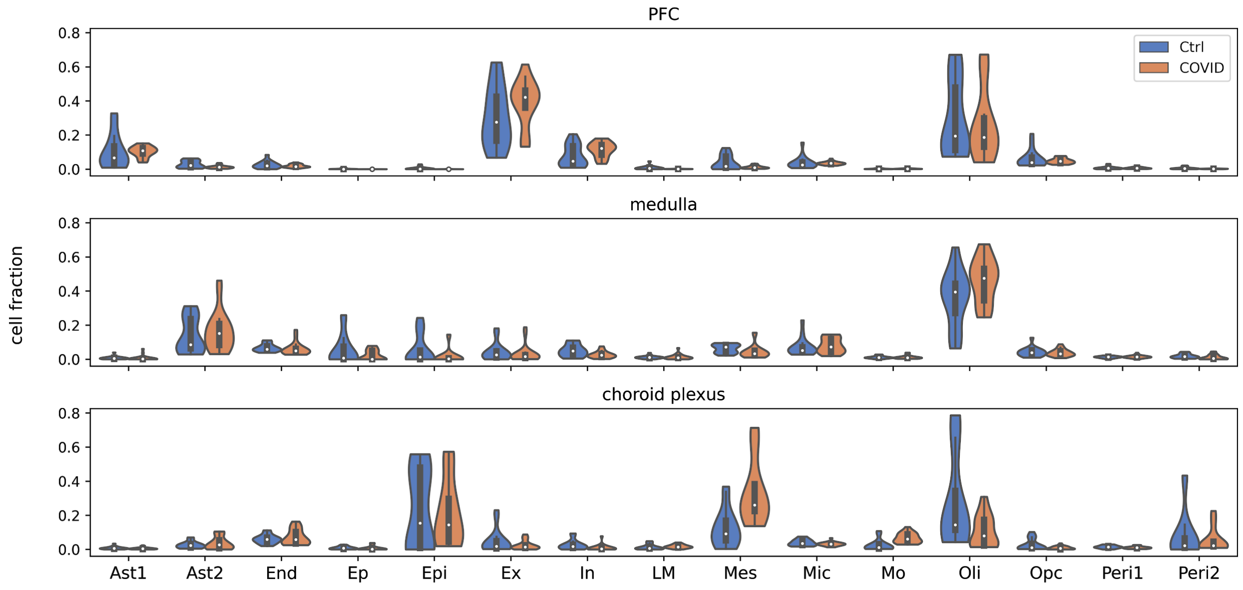


**Fig. S10. Cell type composition among cases and controls.** Visualization is stratified for each brain region: prefrontal cortex (PFC), medulla oblongata and choroid plexus. Orange and blue colors show COVID-19 positive cases and controls, respectively.

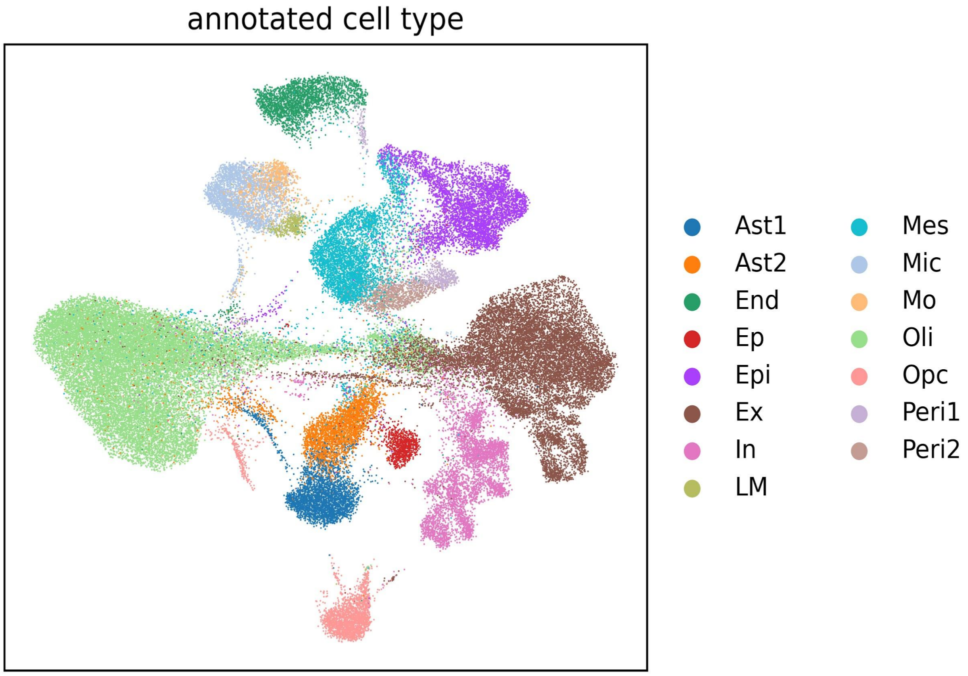


**Fig. S11. UMAP projection based on the activity scores of 131 TF regulons.** Colors show annotated cell types as described in the main text.

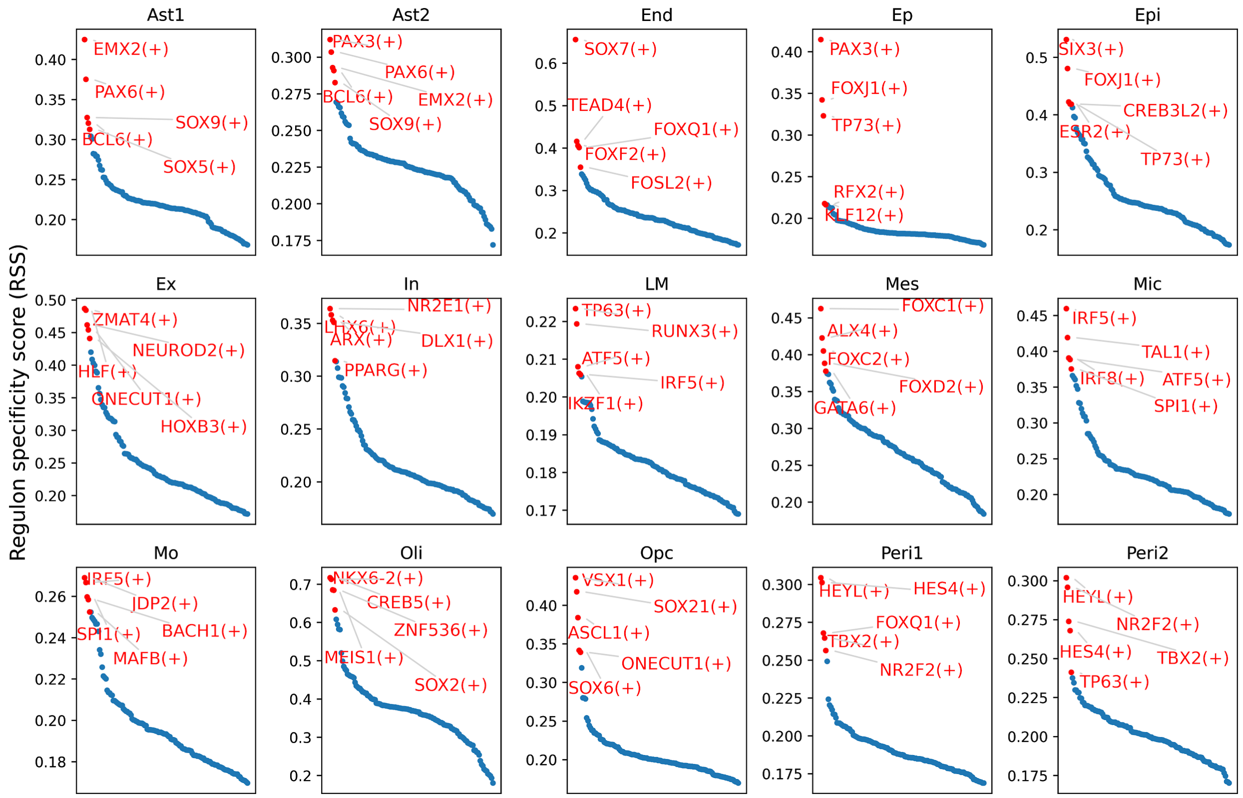


**Fig. S12. Regulon specificity scores (RSS) of each annotated cell population.** A point in a panel shows the RSS of one TF regulon. TF regulons are sorted by the RSSs in each cell type. The top 5 specific regulons are highlighted in red.

**Captions for Data S1 to S8:**

**Data S1. (separate file)**
Clinical characteristics of donors.

**Data S2. (separate file)**
Sequencing statistics for SARS-CoV-2 targeted RNA-seq.

**Data S3. (separate file)**
Gene Ontology term enrichment in differentially expressed genes of PFC microglia.

**Data S4. (separate file)**
Differences in gene set activity scores among COVID-19 patients and controls.

**Data S5. (separate file)**
Genes clustered by pseudo time analysis.

**Data S6. (separate file)**
Transcription regulatory modules estimated by SCENIC.

**Data S7. (separate file)**
Differences in TF regulon activity scores comparing COVID-19 patients and controls.

**Data S8. (separate file)**
Imputed gene associations from COVID-19 GWAS.
